# Describing the fecal metabolome in cryogenically collected samples from healthy participants

**DOI:** 10.1101/708685

**Authors:** Kajetan Trošt, Linda Ahonen, Tommi Suvitaival, Nina Christiansen, Trine Nielsen, Maja Thiele, Suganya Jacobsen, Aleksander Krag, Peter Rossing, Torben Hansen, Lars Ove Dragsted, Cristina Legido-Quigley

## Abstract

**Introduction:** The chemical composition of feces plays an important role in human metabolism. Metabolomics and lipidomics are valuable tools for screening the metabolite composition in feces. Here we set out to describe fecal metabolite composition in healthy participants in frozen stools.

**Methods:** Frozen stool samples were collected from 10 healthy volunteers and cryogenically drilled in four areas along the specimen. Polar metabolites were analyzed using derivatization followed by two-dimensional gas chromatography and time of flight mass spectrometry. Lipids were detected using ultra high-performance liquid chromatography coupled with quadruple time-of-flight mass spectrometry. The technical variation threshold was set to 30% in pooled quality control samples and metabolite variation was then assessed in four areas per specimen. A data-generated network using metabolites found in all areas was computed for healthy participants.

**Results:** 2326 metabolic features were detected. Out of a total of 298 metabolites that were annotated we report here 185 that showed a technical variation of x< 30%. These metabolites included amino acids, fatty acid derivatives, carboxylic acids and phenolic compounds. Lipids predominantly belonged to the groups of diacylglycerols, triacylglycerols and ceramides. Metabolites varied between sampling areas (14%-80%). A network using metabolites present in all areas showed two main clusters, DAG lipids and phenyllactic acid.

**Conclusions:** In feces from healthy participants, the main groups detected were phenolic compounds, ceramides, diacylglycerols and triacylglycerols. Metabolite levels differed considerably depending on the sampling area.

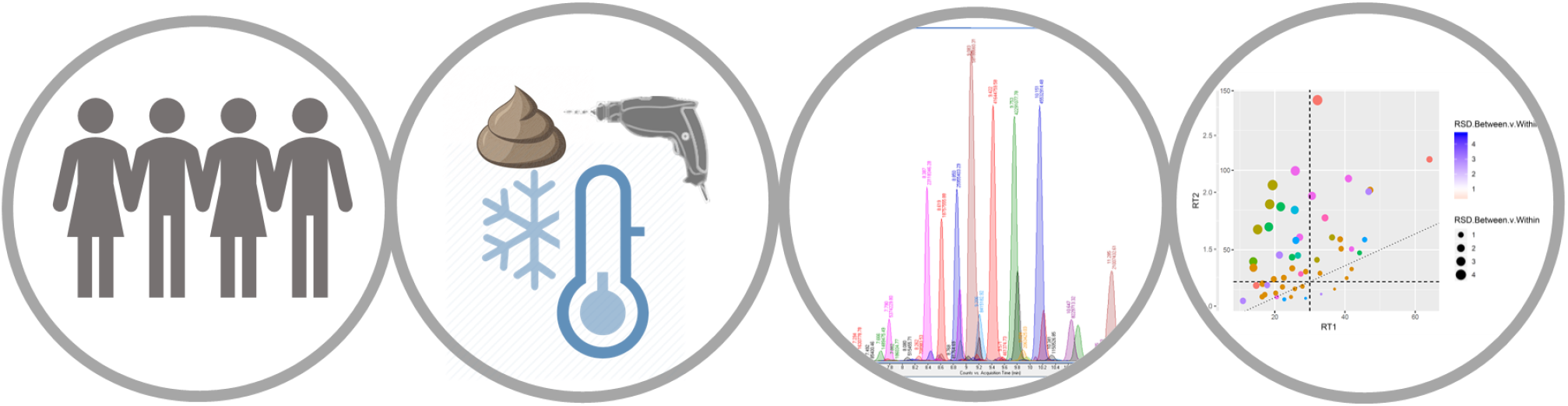

## Introduction

The molecular composition of the intestinal contents is of particular importance in relation to gastrointestinal disorders.^1^ The intestinal microbiota has been associated to metabolic syndrome and its complications (fatty liver disease, obesity and type 2 diabetes)^2,3^ and linked to arthritis, gout, celiac disease and myalgic encephalomyelitis.^4^ The intestinal microbiota affects the metabolism via secondary microbial metabolites and food nutrients^5^. The characterization of the microbiota is traditionally performed via metagenomic profiling.^6^ An emerging approach to determine the influence of microbiota on the host organism is to combine metagenomic profiling with molecular phenotyping. Methods to assess the molecular composition of feces can be targeted and quantitative in concentration or untargeted for explorative purposes.^7^ In contrast to other biofluids (e.g. plasma or urine), stool is highly heterogeneous because it is composed of living bacteria, food remains, nutrients such as lipids, fibers and non-digestible elements.^8,9^ There is little agreement on sample collection techniques and the extent to which a technique can accurately reflect fecal composition.^10–12^ To avoid fermentation, fecal samples can be frozen immediately after collection and stored at −80°C until sample preparation.^13^

Stool is frequently analyzed by targeted analysis focusing on abundant and well characterized metabolites like short chain fatty acids (SCFA) and bile acids. Using gas chromatography (GC), the detection of both volatile and non-volatile metabolites can be achieved.^13^ In order to cover a broader range of metabolites, two-dimensional gas chromatography (GC×GC) can be employed.^14^ These techniques use sample pre-derivatization and can detect many polar compounds, e.g. amino acids, small organic acids, phenols, phenolic acids, certain sugars and medium chain fatty acids. The characterization of lipids is an emerging field that is often studied for metabolic disorders as lipids can make up 15% of the feces composition.^15,16,17^

Several research groups have developed state-of-the-art fecal fingerprinting methods, including chemical labeling with isotopic reagents and high-performance liquid chromatography coupled with high resolution mass spectrometry (HPLC-HRESMS)^18,19^ ^20^, nuclear magnetic resonance (NMR) for compounds like SCFAs and BCAAs^21–23^ and the use of LC-MS^24,25^ and GC-MS^26,27^. In this study, we aim to measure molecules in feces with in-house analytical pipelines using cryogenically drilled samples^28,29,30^, focusing on quality control, sampling procedure and biochemical pathways that can describe feces from healthy volunteers.

## Materials and methods

### Samples

The samples analyzed were from healthy participants, recruited from the general population after a community call for volunteers from Odense University Hospital, under ethical approval from the regional ethics committee of Region of Southern Denmark (S-20160006G). Our inclusion criterium was age 40-75 years. We excluded volunteers in case of: (1) being on any medication, prescription or otherwise, at the time of inclusion, (2) having a chronic disease, whether medicated or not, (3) any use of antibiotics within the six months leading up to inclusion, (4) reported alcohol intake above the low-risk limit of 7 units of alcohol/week for females and 14 units/week for males, or binge drinking (≥5 units at one event). Additionally, volunteers went through a sigmoidoscopy, abdominal ultrasonography and liver elastography to screen for existing gastrointestinal disease. We also performed routine blood tests to rule out diabetes, thyroid disease, dyslipidemia and liver disease. Finally, we checked whether any medical events had occurred during the six months following inclusion. In case of a positive finding, we excluded participants. From February 2016 to December 2016 we included nine men and one female with a mean age of 51 years (range 42-72 years) and a mean BMI of 28 kg/m^2^ (range 25-33). None of the volunteers smoked. One participant had a pacemaker due to arrhythmia, one participant consumed antihistamine for seasonal rhinitis, two consumed a daily vitamin supplement and two ingested fish oil. Volunteers sampled stool in their own home, within 24 hours of a scheduled visit to the research clinic. The volunteer stored the sample in his/her own freezer at −20 °C immediately after sampling. We instructed them to keep the sample on ice, in a cooling bag, during transportation. Upon arrival at the research clinic, we then transferred the stool to a −80 °C storage facility.

For the sampling analyses, each original fecal sample was cryogenically drilled four times at different positions along the specimen. Once the cryogenic drilling was finalized, each drill, 200 mg sample, was mixed with an equal amount of water (50:50, w/w), homogenized and distributed into 20 mg aliquots for further extraction procedures.

The chemicals, sample preparation and instrumental analyses for GC-GC-MS and Lipidomics used in this study are available as *Supplementary information*

### Data pre-processing GC×GC-MS

The detected and potentially identified peaks were aligned using the Gineu software.^32^ The peak count filter was set to 5 in order to filter out features, which did not occur in all samples. Additionally, retention indexes were assigned using the NIST14 and GMD libraries.^33^ The Golm grouping tool was also used for assessment of groups based on characteristic ions. All features that scored less than 850 of similarity score or had more than 35 units of retention index difference were annotated as unknowns. The resulting peak table was exported. Thereafter the data were post-processed and analyzed in R as described later.

### Lipidomics Data Pre-processing

Data processing was performed using MZmine 2.28^38^ using internal mass spectral library and the LIPID MAPS☉ database26. The following steps were applied in the processing: 1) Crop filtering with a m/z range of 200 – 1700 m/z and a RT range of 2.4 to 13.6 min, 2) Mass detection with a noise level of 2500, 3) Chromatogram builder with a min time span of 0.04 min, a min height of 7500 and a m/z tolerance of 0.006 m/z or 10.0 ppm, 4) Chromatogram deconvolution using the local minimum search algorithm with a 70% chromatographic threshold, 0.05 min minimum RT range, 5% minimum relative height, 7500 minimum absolute height, a minimum ratio of peak top/edge of 1 and a peak duration range of 0.04 - 1.0, 5), Isotopic peak grouper with a m/z tolerance of 5.0 ppm, RT tolerance of 0.05 min, maximum charge of 2 and with the most intense isotope set as the representative isotope, 6) Peak filter with minimum 8 data points, a FWHM between 0.0 and 2.0, tailing factor between 0.36 and 2.78 and asymmetry factor between 0.33 and 3.00, 7) Peak list row filter keeping only peak with a minimum of 1 peak in a row, 8) Join aligner with a m/z tolerance of 0.006 or 10.0 ppm and a weight for of 2, a RT tolerance of 0.2 min and a weight of 1 and with no requirement of charge state or ID and no comparison of isotope pattern, 9) Peak list row filter with a minimum of 2 peak in a row, 10) Gap filling using the same RT and m/z range gap filler algorithm with an m/z tolerance of 0.006 m/z or 10.0 ppm, 11) Peak filter with minimum 8 data points, a FWHM between 0.0 and 0.2, tailing factor between 0.36 and 2.78 and asymmetry factor between 0.33 and 3.00 12) Peak list row filter with a minimum of 2 peaks in a row, 13) Identification of lipids using a custom database search with an m/z tolerance of 0.006 m/z or 10.0 ppm and a RT tolerance of 0.2 min, 14) Duplicate peak filter with a m/z tolerance of 0.006 m/z or 10.0 ppm and a RT tolerance of 0.1 min.

### Data post-processing

A schematic figure of the post-processing steps can be seen in **Figure S1**. The data from each of the platforms were post-processed in the same way in R: Exported peak lists were imported to R and each feature was normalized against the most-correlated internal standard in R. Annotated metabolites were assigned with a level 3.^40^ The annotation included features which had equivalent standards injected during the sequence (Level 1) and structure information acquired previously with MS2 fragmentation (Level 2). Annotations were made using our in-house database, the human metabolome database^41^ and LIPIDMAPS^☉39^.

Features at level 4 (unknowns) and with over 30 % coefficient of variation (CV; or, relative standard deviation) in pooled samples were discarded. Further, features with a missing value in more than 20 % of samples were discarded, and remaining missing values were imputed with the k-nearest neighbor algorithm using the impute package.^42^

### Statistical Analysis

Statistical analysis was done in R with the package limma.^43^ Each feature’s variation was compared with a feature-wise F-test, where the feature is the dependent variable and the categorical variable representing the individual is the independent variable. The F-statistic, the associated p-value, and the multiple-testing corrected p-values (Benjamini-Hochberg) were reported.

Coefficients of variation (CV) were computed for each feature as follows, between pooled samples for technical variation, between sampling area for specimen heterogeneity and between participants. These feature-wise value pairs were visualized in two bubble plots for the lipidomics and metabolomics data using the ggplot2 package.^44^ The metabolite/lipid category of each feature was shown by color of the data point to give an overview of the overall variation in the categories. In the integration step, the data were auto-scaled and averaged over replicates of the same individual as well as over features in each compound category. The two resulting individuals-by-compound categories data sets were then combined into a single data matrix, which was visualized as a rows-scaled heatmap using the ggplot2 package to show the relative abundances of each compound category between the individuals.

Data driven partial correlation network of lipids and polar metabolites was computed and visualized with R-package qgraph using the graphical LASSO algorithm and RIC (Rotation Information Criterion). Data were imputed and auto-scaled prior to model-fitting. In the visualization, lipids were shown as circular and polar metabolites as rectangular nodes. Positive and inverse associations between nodes were shown as brown and blue lines, respectively. Strength of the association was shown as width of the line. 3-Phenyl lactic acid was selected as metabolite with most connections and highlighted with purple color. Other nodes were colored by the respective Spearman correlation to 3-Phenyl lactic acid.

## Results and discussion

The metabolome is composed of molecules belonging to a myriad of chemical groups. Untargeted analytical methods are usually tailored to detect a broad chemical group, like lipids or polar metabolites, based on abundance, purity, polarity, volatility and on the availability of bioactive groups for derivatization.^45^ In this study, the preprocessing of the data resulted in the detection of 2326 features in the polar metabolome and lipidome. Among these features, 182 polar and 116 lipid species were putatively identified. Metabolites with relative standard deviations (%RSDs) higher than 30% in the pooled samples were excluded from further statistical evaluation, thus resulting in 185 metabolites fulfilling this requirement **(Figure 1 and 2)**. Annotated features with pool RSD < 30% are presented in **Supplementary Tables S1 and S2** for polar metabolites and lipids respectively

**Figure 1.**
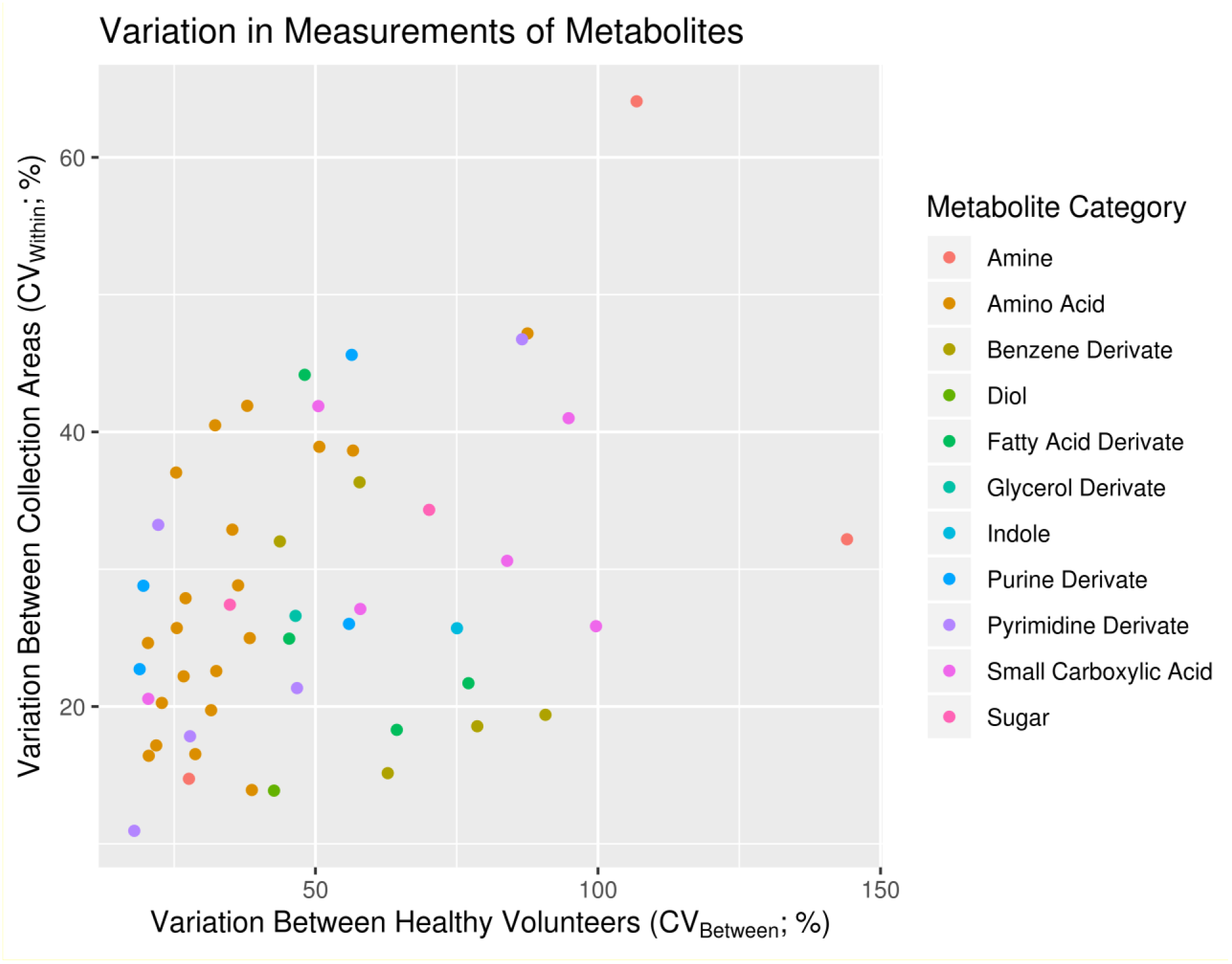
Variation between heathy individuals and sampling area in feces. Each dot represents a metabolite colored by their class. Most metabolites vary more than 30% between collection area and between volunteers making the fecal metabolome highly heterogenous.

**Figure 2.**
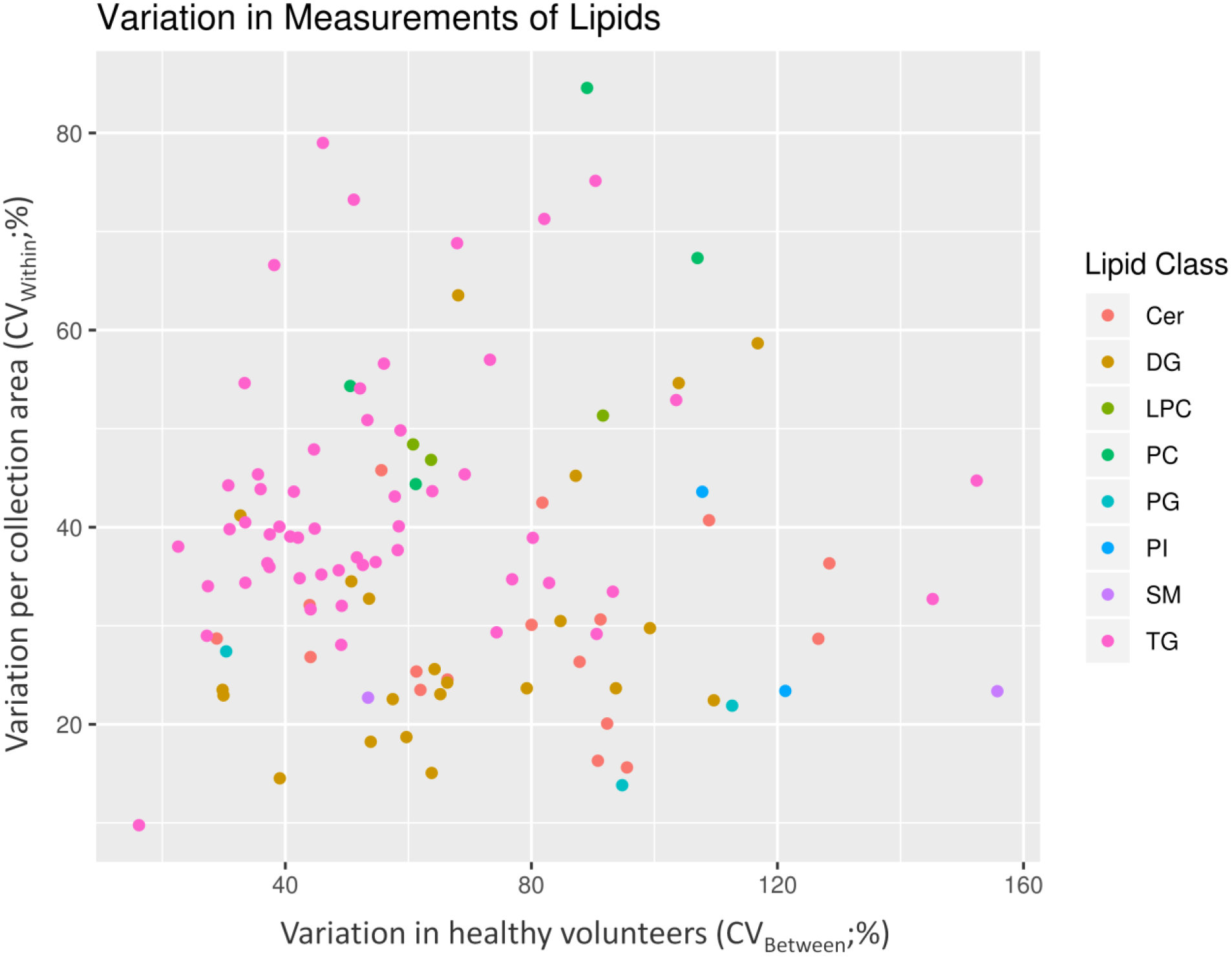
Variation between heathy individuals and sampling area in feces. Each dot represents a lipid colored by their class. Cer: ceramides, DG: diacylglyceride, LPC: lysophosphocholine, PC: phosphatidylcholine, PG: phosphoglycine, PI: phosphoinositol, SM: sphingomyelin, TG triacylglyceride.

To study the feasibility of cryogenically collected samples, we applied additional parameters and calculated metabolite variation in four sampling areas per specimen. As an example of the data collected in this study, the first metabolite in the **Supplementary Tables S1** is 3-Hydroxyphenylacetic acid. The variation between participants for this metabolite exceeded 107% and the variation found in the four sampling area was also high at 64% meaning it is not found homogenously in the stool specimens. Since the technical variation of the measurement is negligible at 4% we can be confident of these results. We then propose that this benzene derivative is found in a wide range of concentrations in the feces of the healthy population. A limitation of this study is that we did not control for the diet of the participants, so the origin of this wide-range normal levels is unknown at this stage. The other benzene derivatives that were detected followed the same trend, apart from 3-Phenyllactic acid, 3-Phenylpropionic acid and 4-Hydroxyphenyllactic acid which were in similar concentrations in the four areas.

### The Fecal metabolome and lipidome

This study allowed for the reporting of 185 molecules, adding to previous research.^47^ These metabolites included amino acids, fatty acid derivates, carboxylic acids, benzene compounds and indole compounds (Figure 1). Fatty acids were mainly present as medium chain fatty acids (e.g. hydroxylated and dicarboxylic species). The group of small carboxylic acids included metabolites from the citric acid cycle as well as hydroxy butyric and propionic acids. Among the amino acids, both regular amino acids as well as branched chain amino acids were present in the feces. In addition to these, some amines, purine derivates and pyrimidine derivates were also annotated. Additionally, the results show that the compounds with the highest variation between participants were benzene derivates (e.g. hydroxyphenyllactic acid, 3-phenyllactic acid and 3-phenylpropionic acid), polyamines and fatty acid derivates (e.g. methylsuccinic acid and 2-hydroxyisocaproic acid), and some of the small carboxylic acids (e.g. tricarballylic acid).

Many of the polar metabolites found in healthy individuals can be associated with bacterial metabolism. A detailed review on microbiota metabolism is out of the scope of this report, but some examples are hydroxyphenyllactic acid which has been found in most anaerobic bacteria and is associated with tyrosine metabolism in humans.^48^ In another study, 3-phenyllactic acid is thought to display antimicrobial properties and to be associated with phenylalanine metabolism.^49^ In addition, the occurrence of 3-phenylpropionic acid is associated with microbially transformed plant polyphenols such as flavanols, flavanones and tannins.^50^ Putrescin is a polyamine produced by the breakdown of amino acids and mainly produced by the microbiota.^51^ Feces are also rich in BCAAs and these can be precursors of hydroxyisocaproic acid, a bacterial end product of leucine degradation.^52^

Clinical studies using the lipidome in fecal samples are sparse in the literature. A study showed its clinical potential in five prematurely born infants.^16^ Van Meulebroek *et al.* also reported the analysis of the fecal lipidomic profile in a small cohort of healthy volunteers and type 2 diabetic patients.^15^ Using adult lyophilized feces, the team identified 127 lipids, out of which 54 were frequently present in feces.

In this study we observed that the fecal samples mainly consisted of ceramides, diglycerides and triglycerides (Figure 2). Lipids, which are commonly detected in plasma (e.g. lysophosphocholines, glycerophosphocholines and sphingomyelins), were also detected in the fecal samples but with remarkably lower coverage.^35^ In fact, only seven phosphatidylcholines could be annotated in the fecal samples while as many as 78 different phosphatidylcholines were identified^35^. Similarly, we detected two sphingomyelins while 30 different sphingomyelins were detected in plasma.^35^

Lipids perform many functions, from storing energy to forming cell and organelle walls or as inflammation markers. Lipids are also a source of nutrition for the bacteria living in the gut, and a fatty diet can be a modifying factor of intestinal microbiota.^53^ Interestingly, the lipid families that have been linked to metabolic syndrome were abundant in the feces of healthy volunteers. Ceramides which are metabolized from sphingomyelins in eukaryotic organisms were abundant in these participants’ feces.^54^ Ceramides are found in large quantities in cell membranes and contribute to cellular signaling mechanisms, particularly cell death, partially reproducing the effect of palmitic acid on insulin signaling.^55^ Triacylglycerols and diacylglycerols are usually thought of as energy metabolites, although they can also be signaling molecules and have long been implicated in the occurrence of metabolic syndrome.^56^

### Correlation between fecal lipids and polar metabolites

Two main clusters were assigned using the partial correlation network inferred with the graphical LASSO algorithm **(Figure 3).** One was composed mostly of diacylglycerides, the other is composed mainly of amino acids and phenyl lactic acid. 3-phenyl lactic acid was the metabolite with most connections to others. Both clusters showed correlations to metabolites from different platforms. Phenyl lactic acid cluster corelated with certain ceramide species and diacylglyceride cluster correlates with a 3-hydroxy butyric acid and 3-phenyl propionic acid.

**Figure 3.**
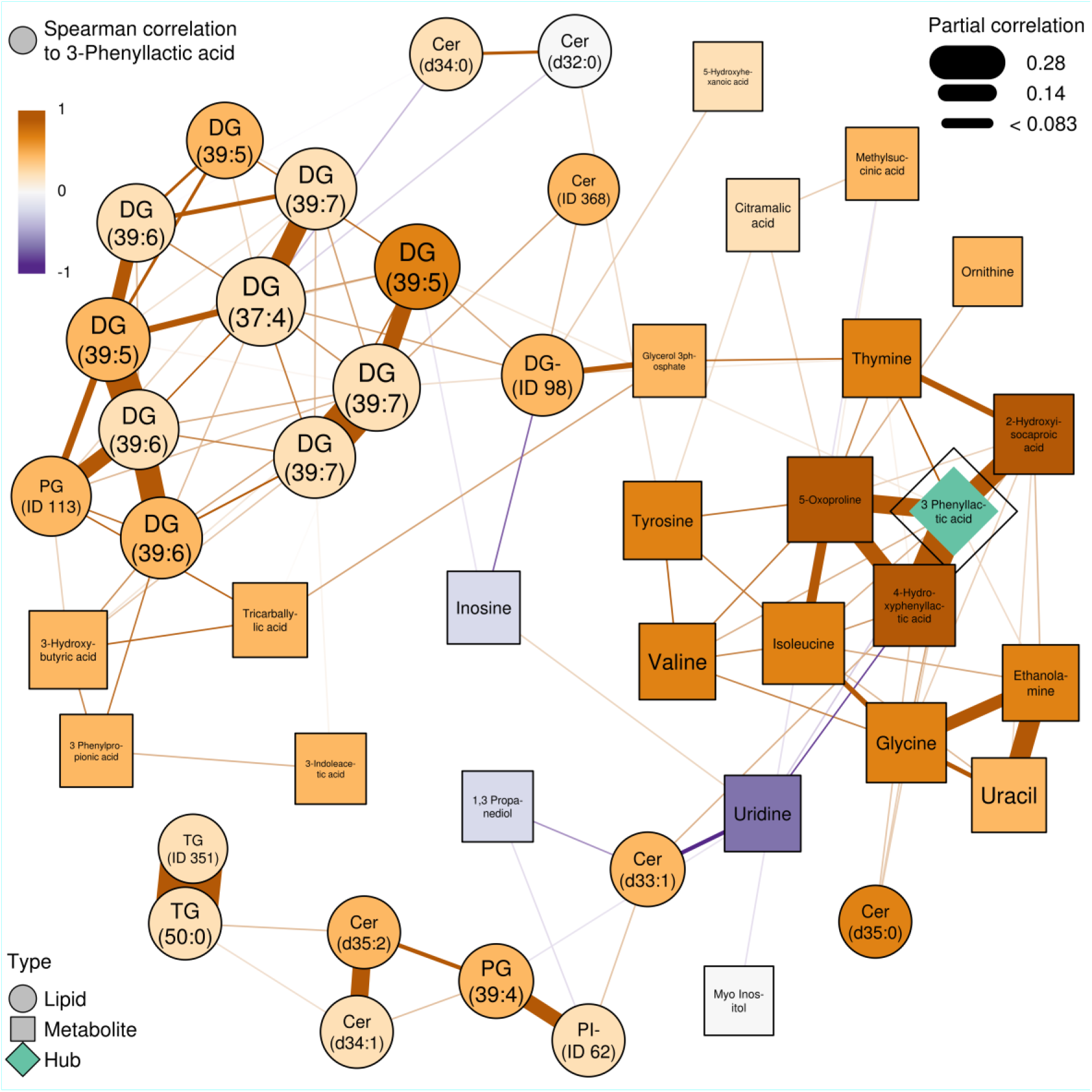
Partial correlation network of fecal polar metabolites and lipids, inferred with the graphical LASSO algorithm. In the figure, lipids were shown as circular and polar metabolites as rectangular nodes. Positive and inverse associations between nodes were shown as brown and purple lines respectively. Strength of the association was shown as width of the line. 3-Phenyl lactic is the main hub of the network with the most associations and is highlighted in green color. Other nodes were colored by the respective Spearman correlation to the hub, 3-Phenyl lactic acid: dark brown is positive correlation and purple is negative correlation. As an example, 4-Hydroxyphenyl lactic acid has a positive partial correlation to 3-Phenyl lactic acid (brown line) as well as a positive Spearman correlation to 3-Phenyl lactic acid (brown node color). The two compounds thus are proportional in abundance in the healthy participants. DGs with the same total number of carbons and double bonds have the same name in the network (ie. 39:7) are different lipids. Cer: ceramides, DG: diacylglyceride, LPC: lysophosphocholine, PC: phosphatidylcholine, PG: phosphoglycine, PI: phosphoinositol, SM: sphingomyelin, TG triacylglyceride.

In the current screening it is not possible to estimate the source of the metabolites. They can derive from food, the microbiota converting food constituents, or they can be primary bacterial or host metabolites. According to the literature, Phenyl lactic acids are mostly associated with lactic acid bacteria and they have antifungal and antimicrobial properties^49^. Additionally, with a focus on integration with metagenomic analysis, the high variability seen in these ten healthy subjects is probably due to variations in both food intake and in microbiota composition^57^. On the other hand, ceramides are known to play a role in metabolic dysfunction^58^. The same is true for diacylglycerides, although their accumulation is thought to be less detrimental than that of triglycerides^59^.

To conclude, the fecal metabolome composition was studied by applying two analytical platforms. The variation of metabolites was assessed for quality control, per area of collection and between healthy participants. Annotated metabolites included a myriad of metabolic pathways, molecules synthesized by the microbiome (phenol derivatives) and three main classes of lipids: ceramides, triacylglycerides and diacylglycerides in the feces of healthy individuals.

## Supporting information

Supporting information to: Describing the fecal metabolome in cryogenically collected samples from healthy participants.

## Acknowledgements

This study has received funding from the European Union’s Horizon 2020 research and innovation programme under grant agreement number 668031 and it was supported by Challenge Grant “MicrobLiver” grant number NNF15OC0016692 from the Novo Nordisk Foundation.

## Supporting information

Methods: Chemicals, sample preparation and instrumental analyses for GC-GC-MS and Lipidomics.

Figure S1: Postprocessing data workflow

Table S1. Selected set of polar metabolites with RSD in pooled samples lower than 30%. Table S2. Selected set of lipids with RSD in pooled samples lower than 30%.

