## Supporting information to: Describing the fecal metabolome in cryogenically collected samples from healthy participants. for "Describing the fecal metabolome in cryogenically collected samples from healthy participants"

**List of supplementary figures and tables:**

Methods: Chemicals, sample preparation and instrumental analyses for GC-GC-MS and Lipidomics.

Figure S1: Postprocessing data workflow

Table S1. Selected set of polar molecules with relative standard deviation lower than 30% in pooled samples.

Table S2. Selected set of lipids with relative standard deviation lower than 30% in pooled samples.

**Methods: Chemicals, sample preparation and instrumental analyses for GC-GC-MS and Lipidomics**

*Chemicals*

Water ($H_2O$), methanol (MeOH), acetonitrile (ACN), isopropanol (IPA) of LC-MS grade were purchased from Honeywell (Morris Plains, NJ, USA). Hexane, sodium chloride (NaCl), formic acid (HCOOH) and chloroform ($CHCl_3$) of reagent grade and ammonium acetate of LC-MS grade were purchased from Sigma-Aldrich (Steinheim, Germany). The derivatization agents methoxyamine hydrochloride (MeOX; TS-45950) and N-methyl-N-trimethylsilyltrifluoroacetamide (MSTFA) were purchased from Thermo Scientific and Sigma-Aldrich, respectively. The group-specific internal standards used for analyzing the polar metabolites were purchased from Sigma-Aldrich and they were as follows: DL-Valine-$d_8$, Heptadecanoic acid-$d_{33}$, Succinic acid-2,2,3,3-$d_4$, L-glutamic acid-2,3,3,4,4-$d_5$, trans Cinnamic acid-$d_7$ and Uric acid-1,3-$^{15}N_2$. Syringe standard 4,4-dibromooctafluorobiphenyl and retention index standards: Undecane (C11), Pentadecane (C15), Heptadecane (C17), Heneicosane (C21) and Pentacosane (C25) were also purchased from Sigma-Aldrich. Quality control of the lipidomics analyses was performed by adding a standard solution containing nine different lipid standards representing the different lipid classes to each sample. The following of these compounds were purchased from Avanti Polar Lipids, Inc. (Alabaster, AL, USA): 1,2-diheptadecanoyl-*sn*-glycero-3-phosphoethanolamine (PE(17:0/17:0)), N-heptadecanoyl-D-*erythro*-sphingosylphosphorylcholine (SM(d18:1/17:0)),N-heptadecanoyl-D-*erythro*-sphingosine (Cer(d18:1/17:0)), 1,2-diheptadecanoyl-*sn*-glycero-3-phosphocholine (PC(17:0/17:0)), 1-heptadecanoyl-2-hydroxy-*sn*-glycero-3-phosphocholine (LPC(17:0)), 1-palmitoyl-d31-2-oleoyl-*sn*-glycero-3-phosphocholine (PC(16:0/d31/18:1)), 1-hexadecyl-2-(9Z-octadecenoyl)-sn-glycero-3-phosphocholine (PC(16:0e/18:1(9Z))), 1-(1Z-octadecenyl)-2-(9Z-octadecenoyl)-sn-glycero-3-phosphocholine (PC(18:0p/18:1(9Z))), 1-octadecanoyl-sn-glycero-3-phosphocholine (LPC(18:0)), 1-(1Z-octadecenyl)-2-docosahexaenoyl-*sn*-glycero-3-phosphocholine (PC(18:0p/22:6)) and 1-stearoyl-2-linoleoyl-*sn*-glycerol (DG(18:0/20:4)). 1,2-dimyristoyl-*sn*-glycero-3-phospho(choline-d13) (PC(14:0/d13)), 1,2,3-triheptadecanoylglycerol (TG(17:0/17:0/17:0)) and 3β-hydroxy-5-cholestene 3-linoleate (ChoE(18:2)) was purchased from Sigma-Aldrich and tripalmitin-1,1,1-13C3 (TG(16:0/16:0/16:0)-13C3), trioctanoin-1,1,1-13C3 (TG(8:0/8:0/8:0)-13C3) and 1-palmitoyl-2-hydroxy-sn-Glycero-3pPhosphatidylcholine (LPC(16:0)) from Larodan AB (Solna, Sweden). Additionally, calibration curves (at concentration levels of 100, 500, 1000, 1500, 2000 and 2500 ng $mL^{-1}$ for the quantification of lipids were prepared using 1-hexadecyl-2-(9Z-octadecenoyl)-sn-glycero-3-phosphocholine (PC(16:0e/18:1(9Z))), 1-(1Z-octadecenyl)-2-(9Z-octadecenoyl)-sn-glycero-3-phosphocholine (PC(18:0p/18:1(9Z))), 1-octadecanoyl-sn-glycero-3-phosphocholine (LPC(18:0)), 1-(1Z-octadecenyl)-2-docosahexaenoyl-*sn*-glycero-3-phosphocholine (PC(18:0p/22:6)), 1-stearoyl-2-arachidonoyl-*sn*-glycero-3-phosphoinositol (PI(18:0/20:4)) and 1-stearoyl-2-linoleoyl-*sn*-glycerol (DG(18:0/18:2)) from Avanti Polar Lipids, Inc., 1-Palmitoyl-2-Hydroxy-sn-Glycero-3-Phosphatidylcholine (LPC(16:0)) from Larodan, and 1,2,3-Triheptadecanoylglycerol (TG(17:0/17:0/17:0)) and 3β-Hydroxy-5-cholestene 3-linoleate (ChoE(18:2)), 3β-Hydroxy-5-cholestene 3-oleate (ChoE(18:1(9Z))), 5-Cholesten-3β-yl octadecenoate (ChoE(18:0)) and 5-Cholestene 3-palmitate (ChoE(16:0)) from Sigma-Aldrich.

*Sample preparation*

The method used for the analysis of polar metabolites was originally developed for the analysis of plasma and cerebrospinal fluid[3] and modified to fit the specific matrix (i.e. fecal samples) and the target analytes. An internal standard mixture consisting of 25 mg $L^{-1}$ DL-Valine-$d_8$, 160 mg $L^{-1}$ Heptadecanoic acid-$d_{33}$, 25 mg $L^{-1}$ Succinic acid-$d_4$, 100 mg $L_{-1}$ L-glutamic acid-$d_5$, 100 mg $L^{-1}$ trans-Cinnamic acid-$d_7$ and 100 mg $L^{-1}$ Uric acid-$^{15}N_2$ was created by dissolving the analytes in methanol. After this, 20mg of each homogenized sample was mixed with 600 µL of $H_2O$ and 60 µL of the internal standard mixture. Samples were vortex-mixed and sonicated for 5 minutes after which they were incubated on ice for 30 min and centrifuged (9400 × g, 5 min, 4 °C). The supernatants were filtered through Millipore Millex PVDF syringe filters with diameter of 4 mm and pore size of 0.45 µm. Finally, 150 µL of the filtered extracts were transferred to glass vials and evaporated to dryness before further analysis. The samples were derivatized using a previously described MeOX:MSTFA procedure.[31] Here, the derivatization converts the reactive biological groups into trimethylsilyl derivates, which increase the volatility of the biomolecules. The derivatization was performed automatically using a MultiPurpose Sampler 2 (MPS2, Gerstel; Mülheim an der Ruhr, Germany) with two robotic hands. First, 25 µL of MeOX was added to each sample after which they were incubated and shaked for one hour at 45°C. Then, 25 µL of MSTFA (including a set of n-alkanes as retention index standards at 8 mg $L^{-1}$) was added to each sample and the samples were again incubated and shaked at 45 °C for one hour. Finally, before injecting the sample for analysis, 50 µL of the injection standard 4,4′-dibromooctafluorobiphenyl (9.8 mg $L^{-1}$ in hexane) was added to each sample.

*Analysis of metabolites by GC×GC-MS*

*Sample analysis*

The polar metabolites were analyzed using a Pegasus 4D (LECO; Saint Joseph; USA) system, which combines two-dimensional chromatographic separation with time-of-flight (TOF) mass spectrometric detection. A volume of 1 µL of the derivatized sample extract was injected into the chromatographic system Chromatographic separation was then achieved using a system of two columns guarded by a retention gap column (1.7m, 0.53 mm ID, FS deactivated) from Agilent Technologies (Santa Clara, CA, USA). The primary separation was achieved using a 10 m × 0.18 mm I.D. Rxi-5 ms (Restek Corp., Bellefonte, PA, USA) column and the secondary column was a 1.5 m × 0.1 mm I.D. BPX-50 (SGE Analytical Science, Austin, TX, USA). The temperature gradient used for separating the analytes was as follows: 50 °C (2 min), 7°C/min to 240°C, 25°/min to 300 °C (3 min). The secondary oven temperature was set to be 20 °C higher than primary oven temperature. The modulator cycle was set to 4s, producing narrow chromatographic peaks of approximately 0.2s. Hence, the speed of the detector was set to 100 Hz. The ChromaTOF software (version 4.32; LECO Corporation, St. Joseph, USA) was used for all data acquisition as well as for processing of the raw data. The default peak picking criteria for the raw data processing was set to 15s in the first dimension and 0.2 in second dimension and the signal to noise ratio was set to 100. The NIST14 spectral library was used for potential identification of metabolites and the created text files were exported for the further data processing.

*Lipidomics analyses*

*Sample preparation*

Previously published lipidomics procedures,[34-36] which have been validated for plasma samples, were used for the fecal lipidomics with some modifications. The 40 drilled fecal samples (50 mg of fecal slurry) were prepared using a modified Folch-extraction procedure.[37] To ensure high quality of data, blank samples and a pool of all fecal samples were prepared using the same protocol and they were analyzed along with the actual samples. Briefly, 10 µL of 0.9% NaCl, 92 µL of CHCl3:MeOH (2:1, v/v) and 28 µL of a 10 µg/mL internal standard solution (containing PE(17:0/17:0), SM(d18:1/17:0), Cer(d18:1/17:0), PC(17:0/17:0), LPC(17:0), PC(16:0/d31/18:1), PC(14:0/d13), TG(16:0/16:0/16:0)-13C3 and TG(8:0/8:0/8:0)-13C3) were added to each fecal sample. The samples were vortex mixed and incubated on ice for 30 min after which they were centrifuged (9400 × *g*, 3 min, 4 °C). Finally, 60 µL from the lower layer of each sample was then transferred to a glass vial with an insert and 60 µL of $CHCl_3$:MeOH (2:1, v/v) was added to each sample. All samples were stored in -80 °C until analysis.

*Sample analysis*

The samples were analyzed using an UHPLC-Q-TOF-MS, which has been presented in detail previously.[36] Briefly, the UHPLC system was a 1290 Infinity system from Agilent Technologies. The system was equipped with a multisampler (maintained at 10 °C), a quaternary solvent manager and a column thermostat (maintained at 50 °C). Separations were performed on an ACQUITY UPLC® BEH C18 column (2.1 mm × 100 mm, particle size 1.7 µm) by Waters (Milford, USA). The flow rate was 0.4 mL $min^{-1}$ and the injection volume was 1 µL. $H_2O$ + 1% $NH_4Ac$ (1M) + 0.1% HCOOH (A) and ACN:IPA (1:1, v/v) + 1% $NH_4Ac$ + 0.1% HCOOH (B) were used as the mobile phases for gradient elution of the analytes and the gradient was as follows: from 0 to 2 min 35-80% B, from 2 to 7 min 80-100% B and from 7 to 14 min 100% B.

The mass spectrometer coupled to the UHPLC system was a 6550 iFunnel quadrupole time of flight (Q-TOF) from Agilent Technologies. The Q-TOF was interfaced with a dual jet stream electrospray (dual ESI) ion source. Nitrogen generated by a nitrogen generator (PEAK Scientific, Scotland, UK) was used as the nebulizing gas at a pressure of 21 psi, as the drying gas at a flow rate of 14 L $min^{-1}$ (at 193 °C) and as the sheath gas at a flow rate of 11 L $min^{-1}$ (at 379 °C). Pure nitrogen (6.0) from Strandmøllen A/S (Klampenborg, Denmark) was used as the collision gas. The capillary voltage and the nozzle voltage were kept at 3643 and 1500 V, respectively. The reference mass solution including ions at m/z 121.0509 and 922.0098 was prepared according to instructions by Agilent and it was introduced to the mass spectrometer through the other nebulizer in the dual ESI ion source using a separate Agilent series 1290 isocratic pump at a constant flow rate of 4 mL/min (split to 1:100). The acquisition mass range was m/z 100–1700 and the instrument was run using the extended dynamic range with an approximate resolution of 30 000 FWHM measured at m/z 1521.9715. MassHunters B.06.01 (Agilent Technologies) software was used for all data acquisition.

**Supplementary Figure S1: Postprocessing data workflow**

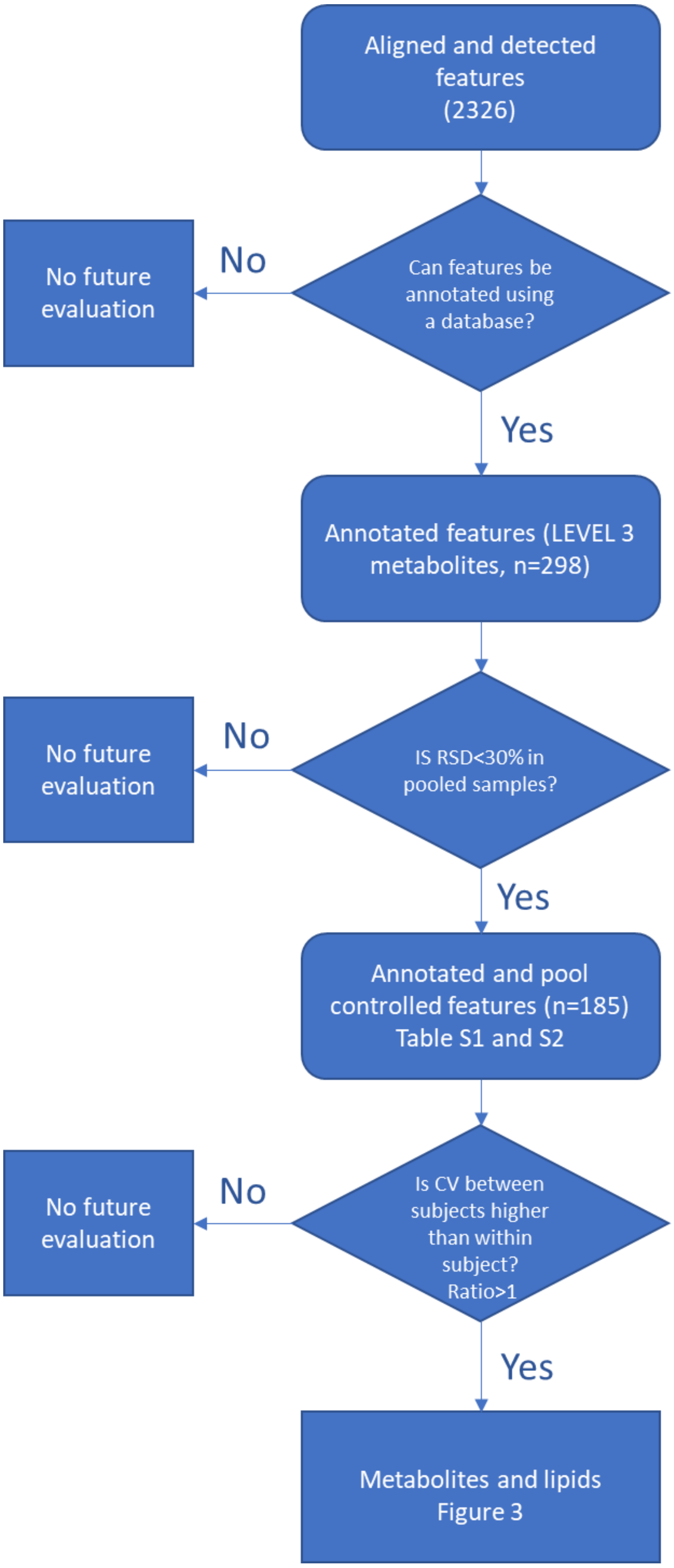

**Supplementary Table S1. Selected set of polar molecules with relative standard deviation lower than 30% in pooled samples.**

| | Name | CV - pooled sample (%) | CV Between Subjects (%) | CV Within Drills (%) | Ratio (CV between /CV within) | feature-wise F-test | adj.P.Val |
|---|---|---|---|---|---|---|---|
| **BENZENE DERIVATES** | | | | | | | |
| | 3-Hydroxyphenylacetic acid | 4 | 107 | 68 | 1.6 | 6 | 8.26E-05 |
| | Tyramine | 8 | 98 | 44 | 2.2 | 18 | 7.12E-10 |
| | 3-Phenyllactic acid | 9 | 79 | 19 | 4.2 | 112 | 3.47E-20 |
| | 3-Phenylpropionic acid | 10 | 63 | 15 | 4.2 | 215 | 1.19E-23 |
| | 4-Hydroxyphenyllactic acid | 12 | 91 | 19 | 4.7 | 142 | 1.53E-21 |
| | 3-Hydroxyphenylpropionic acid | 12 | 58 | 36 | 1.6 | 32 | 7.31E-13 |
| | 4-Hydroxybenzeneacetic acid | 13 | 45 | 64 | 0.7 | 7 | 9.19E-06 |
| | Phtalic acid | 20 | 92 | 38 | 2.4 | 16 | 2.51E-09 |
| | Benzeneacetic acid | 24 | 44 | 32 | 1.4 | 24 | 2.34E-11 |
| | 3,4-Dihydroxyhydrocinnamic acid | 24 | 95 | 52 | 1.9 | 7 | 1.36E-05 |
| **INDOLS** | | | | | | | |
| | 3-Indoleacetic acid | 20 | 75 | 26 | 2.9 | 51 | 1.22E-15 |
| | 3-Indolepropionic acid | 23 | 74 | 45 | 1.6 | 9 | 1.59E-06 |
| **CARBOXYLIC ACIDS** | | | | | | | |
| | Fumaric acid | 8 | 77 | 67 | 1.2 | 8 | 3.73E-06 |
| | Maleic acid | 10 | 20 | 21 | 1.0 | 67 | 3.73E-17 |
| | Lactic Acid | 11 | 66 | 71 | 0.9 | 8 | 3.66E-06 |
| | 2-Hydroxybutyric acid | 14 | 84 | 31 | 2.7 | 92 | 4.54E-19 |
| | Citric acid | 15 | 51 | 37 | 1.4 | 13 | 2.79E-08 |
| | Citramalic acid | 22 | 46 | 30 | 1.6 | 12 | 7.95E-08 |
| | 3-Hydroxybutyric acid | 25 | 58 | 27 | 2.1 | 83 | 2.10E-18 |
| | Tricarballylic acid | 28 | 100 | 26 | 3.9 | 25 | 1.37E-11 |
| | Succinic acid | 30 | 40 | 49 | 0.8 | 15 | 7.95E-09 |
| | 2,4-Dihydroxybutyric acid | 30 | 58 | 26 | 2.2 | 11 | 1.35E-07 |
| **FATTY ACID DERIVATES** | | | | | | | |
| | Pentadecanoic acid | 3 | 48 | 44 | 1.1 | 21 | 1.14E-10 |
| | Palmitic Acid | 3 | 33 | 49 | 0.7 | 12 | 1.20E-07 |
| | Azelaic acid | 4 | 39 | 66 | 0.6 | 10 | 3.53E-07 |
| | 3-Hydroxyisovaleric acid | 6 | 54 | 33 | 1.6 | 6 | 6.42E-05 |
| | 5-Hydroxyhexanoic acid | 7 | 45 | 25 | 1.8 | 51 | 1.22E-15 |
| | Stearic acid | 9 | 53 | 44 | 1.2 | 18 | 5.93E-10 |
| | Suberic acid | 9 | 51 | 67 | 0.8 | 9 | 1.66E-06 |
| | Arachidic acid | 12 | 34 | 54 | 0.6 | 9 | 1.69E-06 |
| | Methylsuccinic acid | 14 | 64 | 18 | 3.5 | 201 | 2.11E-23 |
| | 2-Hydroxyisocaproic acid | 15 | 77 | 22 | 3.6 | 65 | 4.63E-17 |
| | Linolenic acid | 15 | 144 | 83 | 1.7 | 9 | 2.33E-06 |
| | Hexanoic acid | 17 | 89 | 68 | 1.3 | 7 | 1.22E-05 |
| | Sebacic acid | 18 | 52 | 50 | 1.1 | 13 | 3.29E-08 |
| | Nonanoic acid | 19 | 69 | 62 | 1.1 | 8 | 6.00E-06 |
| | Dodecanoic acid | 28 | 182 | 80 | 2.3 | 4 | 0.00279 |
| | Decanoic acid | 30 | 86 | 63 | 1.4 | 5 | 0.000263 |
| | Myristic acid | 30 | 87 | 70 | 1.2 | 6 | 9.34E-05 |

| | | | | | | | |
|---|---|---|---|---|---|---|---|
| **AMINOACIDS** | | | | | | | |
| | Isoleucine | 6 | 27 | 22 | 1.2 | 79 | 3.53E-18 |
| | 5-Oxoproline | 7 | 29 | 17 | 1.7 | 150 | 1.02E-21 |
| | Leucine | 7 | 20 | 25 | 0.8 | 64 | 5.21E-17 |
| | Glycine_iso1 | 8 | 21 | 16 | 1.3 | 121 | 1.21E-20 |
| | Alanine | 9 | 39 | 14 | 2.8 | 148 | 1.07E-21 |
| | Tyrosine_iso1 | 10 | 22 | 17 | 1.3 | 144 | 1.42E-21 |
| | Tyrosine_iso2 | 12 | 36 | 29 | 1.3 | 41 | 2.55E-14 |
| | Acetyl-Lysine | 13 | 26 | 26 | 1.0 | 45 | 6.19E-15 |
| | Ornithine | 13 | 32 | 20 | 1.6 | 104 | 8.64E-20 |
| | Valine | 13 | 23 | 20 | 1.1 | 102 | 1.21E-19 |
| | Aspartic acid_iso1 | 16 | 33 | 23 | 1.4 | 80 | 3.00E-18 |
| | Threonine_iso1 | 16 | 27 | 28 | 1.0 | 47 | 3.76E-15 |
| | Serine_iso1 | 18 | 66 | 65 | 1.0 | 9 | 2.28E-06 |
| | Threonine_iso2 | 19 | 60 | 68 | 0.9 | 8 | 4.53E-06 |
| | Tryptophan_iso1 | 20 | 35 | 33 | 1.1 | 32 | 6.46E-13 |
| | Methionine | 20 | 65 | 64 | 1.0 | 9 | 1.83E-06 |
| | Phenylalanine_iso1 | 20 | 67 | 63 | 1.1 | 9 | 2.55E-06 |
| | Aspartic acid_iso2 | 22 | 41 | 43 | 0.9 | 19 | 3.12E-10 |
| | Tryptophan_iso2 | 22 | 51 | 39 | 1.3 | 21 | 1.42E-10 |
| | Serine_iso2 | 22 | 25 | 37 | 0.7 | 28 | 3.10E-12 |
| | Proline | 22 | 38 | 42 | 0.9 | 26 | 1.10E-11 |
| | Asparagine | 23 | 122 | 56 | 2.2 | 8 | 2.85E-06 |
| | Glycine_iso2 | 23 | 38 | 25 | 1.5 | 69 | 2.66E-17 |
| | Phenylalanine_iso2 | 24 | 25 | 44 | 0.6 | 19 | 3.30E-10 |
| | 4-Aminobutanoic acid | 27 | 65 | 68 | 1.0 | 9 | 1.59E-06 |
| **AMINES** | | | | | | | |
| | Ethanolamine_iso_1 | 7 | 28 | 15 | 1.9 | 129 | 4.80E-21 |
| | Putrescine | 22 | 144 | 32 | 4.5 | 65 | 4.96E-17 |
| | Ethanolamine_iso_2 | 26 | 31 | 31 | 1.0 | 20 | 2.73E-10 |
| | Cadaverine | 30 | 107 | 64 | 1.7 | 27 | 6.09E-12 |
| **PURINE DERIVATES** | | | | | | | |
| | Inosine | 15 | 56 | 26 | 2.2 | 50 | 1.52E-15 |
| | Hypoxanthine | 20 | 19 | 23 | 0.8 | 65 | 4.63E-17 |
| | 2-Deoxyinosine | 28 | 56 | 46 | 1.2 | 20 | 2.05E-10 |
| **PYRIMIDINE DERIVATES** | | | | | | | |
| | Uracil | 7 | 18 | 11 | 1.6 | 290 | 2.65E-25 |
| | Thymine | 7 | 28 | 18 | 1.6 | 132 | 3.95E-21 |
| | Orotic Acid | 8 | 87 | 47 | 1.9 | 50 | 1.61E-15 |
| | Uridine_iso1 | 25 | 47 | 21 | 2.2 | 95 | 3.24E-19 |
| | Uridine_iso2 | 29 | 41 | 39 | 1.1 | 20 | 2.70E-10 |
| **DIOLS** | | | | | | | |
| | 1,3-Propanediol | 11 | 43 | 14 | 3.1 | 164 | 3.52E-22 |
| **GLYCEROL DERIVATES** | | | | | | | |
| | Glycerol-3-phosphate | 30 | 47 | 27 | 1.8 | 56 | 4.08E-16 |
| **STEROIDS** | | | | | | | |
| | Cholesterol | 4 | 142 | 54 | 2.7 | 19 | 3.34E-10 |
| **SUGARS** | | | | | | | |
| | Myo-Inositol | 8 | 35 | 27 | 1.3 | 47 | 3.76E-15 |
| | Cellobiose | 22 | 70 | 34 | 2.0 | 37 | 9.52E-14 |

**Supplementary Table S2. Selected set of lipids with relative standard deviation lower than 30% in pooled samples.**

| ID | Name | CV - pooled sample (%) | CV Between Subjects (%) | CV Within Drills (%) | Ratio (CV between /CV within) | feature-wise F-test | adj.P.Val |
|---|---|---|---|---|---|---|---|
| **DIACYLGLYCEROL LIPIDS** | | | | | | | |
| | DG(23:0) | 8 | 87 | 45 | 1.9 | 34 | 1.74E-13 |
| | DG(32:1) | 3 | 64 | 26 | 2.5 | 87 | 5.08E-19 |
| | DG(33:09) | 15 | 85 | 31 | 2.8 | 44 | 5.70E-15 |
| | DG(36:3) | 12 | 68 | 64 | 1.1 | 6 | 9.12E-05 |
| | DG(36:4) | 8 | 104 | 55 | 1.9 | 8 | 4.34E-06 |
| | DG(37:1) | 10 | 51 | 35 | 1.5 | 7 | 2.02E-05 |
| | DG(37:4) | 3 | 79 | 24 | 3.4 | 41 | 1.60E-14 |
| | DG(39:5) | 12 | 60 | 19 | 3.2 | 39 | 2.42E-14 |
| | DG(39:5) | 6 | 54 | 18 | 3.0 | 44 | 6.43E-15 |
| | DG(39:5) | 2 | 58 | 23 | 2.6 | 76 | 3.21E-18 |
| | DG(39:6) | 13 | 64 | 15 | 4.2 | 102 | 6.28E-20 |
| | DG(39:6) | 3 | 39 | 15 | 2.7 | 56 | 2.04E-16 |
| | DG(39:6) | 12 | 66 | 24 | 2.7 | 102 | 6.37E-20 |
| | DG(39:6) | 7 | 54 | 33 | 1.6 | 41 | 1.39E-14 |
| | DG(39:7) | 4 | 110 | 22 | 4.9 | 133 | 2.33E-21 |
| | DG(39:7) | 4 | 65 | 23 | 2.8 | 81 | 1.23E-18 |
| | DG(39:7) | 5 | 99 | 30 | 3.3 | 94 | 1.89E-19 |
| | DG(ID_109) | 4 | 30 | 23 | 1.3 | 60 | 9.46E-17 |
| | DG(ID_155) | 12 | 117 | 59 | 2.0 | 8 | 3.49E-06 |
| | DG(ID_206) | 3 | 33 | 41 | 0.8 | 22 | 4.71E-11 |
| | DG(ID_90) | 4 | 30 | 24 | 1.3 | 43 | 9.02E-15 |
| | DG(ID_98) | 15 | 94 | 24 | 4.0 | 42 | 1.16E-14 |
| **TRIACYLGLYCEROL LIPIDS** | | | | | | | |
| | TG(45:0) | 23 | 38 | 67 | 0.6 | 7 | 8.82E-06 |
| | TG(46:0) | 15 | 80 | 39 | 2.1 | 28 | 2.10E-12 |
| | TG(48:1) | 10 | 93 | 34 | 2.8 | 6 | 8.44E-05 |
| | TG(49:3) | 1 | 16 | 10 | 1.7 | 70 | 9.49E-18 |
| | TG(50:0) | 19 | 44 | 32 | 1.4 | 29 | 1.32E-12 |
| | TG(50:0) | 16 | 91 | 29 | 3.1 | 149 | 6.01E-22 |
| | TG(50:1) | 16 | 49 | 32 | 1.5 | 26 | 4.79E-12 |
| | TG(50:1) | 13 | 58 | 40 | 1.5 | 13 | 1.79E-08 |
| | TG(50:2) | 10 | 53 | 36 | 1.5 | 18 | 4.82E-10 |
| | TG(50:3) | 12 | 34 | 41 | 0.8 | 28 | 2.10E-12 |
| | TG(50:4) | 18 | 59 | 50 | 1.2 | 10 | 4.39E-07 |
| | TG(51:1) | 14 | 145 | 33 | 4.4 | 25 | 7.02E-12 |
| | TG(51:1) | 13 | 152 | 45 | 3.4 | 31 | 5.09E-13 |
| | TG(51:2) | 16 | 68 | 69 | 1.0 | 7 | 2.86E-05 |
| | TG(51:3) | 24 | 33 | 55 | 0.6 | 12 | 7.00E-08 |
| | TG(51:4) | 20 | 73 | 57 | 1.3 | 5 | 0.000245 |
| | TG(52:1) | 8 | 58 | 43 | 1.3 | 18 | 4.73E-10 |
| | TG(52:1) | 18 | 45 | 48 | 0.9 | 19 | 1.93E-10 |
| | TG(52:2) | 14 | 42 | 39 | 1.1 | 13 | 1.88E-08 |

| | | | | | | | |
|---|---|---|---|---|---|---|---|
| | TG(52:2) | 16 | 39 | 40 | 1.0 | 21 | 5.44E-11 |
| | TG(52:3) | 5 | 49 | 36 | 1.4 | 37 | 5.86E-14 |
| | TG(52:3) | 13 | 41 | 39 | 1.0 | 7 | 8.49E-06 |
| | TG(52:4) | 4 | 46 | 35 | 1.3 | 30 | 8.45E-13 |
| | TG(52:4) | 5 | 27 | 34 | 0.8 | 37 | 5.95E-14 |
| | TG(52:5) | 5 | 23 | 38 | 0.6 | 26 | 4.15E-12 |
| | TG(52:6) | 2 | 82 | 71 | 1.2 | 10 | 5.06E-07 |
| | TG(54:3) | 16 | 31 | 40 | 0.8 | 25 | 7.11E-12 |
| | TG(54:3) | 6 | 37 | 36 | 1.0 | 16 | 2.07E-09 |
| | TG(54:4) | 8 | 52 | 54 | 1.0 | 4 | 0.00182 |
| | TG(54:4) | 8 | 31 | 44 | 0.7 | 19 | 1.94E-10 |
| | TG(54:5) | 5 | 34 | 34 | 1.0 | 30 | 7.72E-13 |
| | TG(54:6) | 1 | 37 | 36 | 1.0 | 42 | 1.04E-14 |
| | TG(54:6) | 0 | 41 | 44 | 0.9 | 19 | 2.41E-10 |
| | TG(56:4) | 3 | 69 | 45 | 1.5 | 23 | 2.04E-11 |
| | TG(56:5) | 6 | 42 | 35 | 1.2 | 29 | 1.44E-12 |
| | TG(58:4) | 3 | 90 | 75 | 1.2 | 5 | 0.000211 |
| | TG(ID_105) | 13 | 53 | 51 | 1.1 | 23 | 2.13E-11 |
| | TG(ID_144) | 14 | 36 | 45 | 0.8 | 25 | 7.11E-12 |
| | TG(ID_146) | 23 | 77 | 35 | 2.2 | 30 | 7.31E-13 |
| | TG(ID_159) | 21 | 46 | 79 | 0.6 | 9 | 1.38E-06 |
| | TG(ID_166) | 20 | 51 | 73 | 0.7 | 3 | 0.00482 |
| | TG(ID_192) | 13 | 55 | 37 | 1.5 | 33 | 3.02E-13 |
| | TG(ID_198) | 12 | 58 | 38 | 1.6 | 8 | 4.50E-06 |
| | TG(ID_220) | 12 | 104 | 53 | 2.0 | 8 | 6.30E-06 |
| | TG(ID_246) | 20 | 36 | 44 | 0.8 | 25 | 7.42E-12 |
| | TG(ID_308) | 7 | 38 | 39 | 1.0 | 9 | 1.11E-06 |
| | TG(ID_315) | 8 | 27 | 29 | 0.9 | 41 | 1.39E-14 |
| | TG(ID_351) | 18 | 74 | 29 | 2.5 | 52 | 7.37E-16 |
| | TG(ID_57) | 3 | 52 | 37 | 1.4 | 4 | 0.000791 |
| | TG(ID_94) | 14 | 64 | 44 | 1.5 | 4 | 0.000877 |
| | TG(ID_99) | 11 | 83 | 34 | 2.4 | 2 | 0.0271 |
| **CERAMIDES** | | | | | | | |
| | Cer(d27:1) | 8 | 128 | 36 | 3.5 | 2 | 0.142 |
| | Cer(d29:1) | 8 | 82 | 43 | 1.9 | 2 | 0.0392 |
| | Cer(d32:0) | 14 | 96 | 16 | 6.1 | 110 | 2.49E-20 |
| | Cer(d33:1) | 23 | 62 | 24 | 2.6 | 35 | 1.04E-13 |
| | Cer(d34:0) | 15 | 66 | 25 | 2.7 | 30 | 7.76E-13 |
| | Cer(d34:1) | 25 | 61 | 25 | 2.4 | 105 | 4.45E-20 |
| | Cer(d34:1) | 5 | 88 | 26 | 3.3 | 134 | 2.32E-21 |
| | Cer(d35:0) | 14 | 127 | 29 | 4.4 | 124 | 4.80E-21 |
| | Cer(d35:1) | 15 | 29 | 29 | 1.0 | 12 | 6.94E-08 |
| | Cer(d35:1) | 17 | 44 | 32 | 1.4 | 31 | 4.94E-13 |
| | Cer(d35:2) | 22 | 91 | 16 | 5.6 | 241 | 1.98E-24 |
| | Cer(d36:0) | 11 | 92 | 20 | 4.6 | 149 | 6.01E-22 |
| | Cer(d40:2) | 6 | 109 | 41 | 2.7 | 87 | 5.08E-19 |
| | Cer(d42:2) | 5 | 91 | 31 | 3.0 | 127 | 3.79E-21 |
| | Cer(d42:2) | 5 | 80 | 30 | 2.7 | 96 | 1.47E-19 |
| | Cer(ID_347) | 8 | 56 | 46 | 1.2 | 8 | 4.97E-06 |
| | Cer(ID_368) | 8 | 44 | 27 | 1.6 | 30 | 9.34E-13 |
| **LYSOPHOSPHATIDYLCHOLINES** | | | | | | | |
| | LPC(16:0) | 18 | 64 | 47 | 1.4 | 11 | 1.68E-07 |

| | | | | | | | |
|---|---|---|---|---|---|---|---|
| | LPC(18:0) | 7 | 61 | 48 | 1.3 | 22 | 4.71E-11 |
| | LPC(18:1) | 19 | 92 | 51 | 1.8 | 5 | 0.00042 |
| **GLYCEROPHOSPHOCHOLINES** | | | | | | | |
| | PC(34:1) | 8 | 61 | 44 | 1.4 | 13 | 2.31E-08 |
| | PC(34:2) | 11 | 51 | 54 | 0.9 | 5 | 0.000365 |
| | PC(36:3) | 9 | 107 | 67 | 1.6 | 4 | 0.00125 |
| | PC(36:4) | 13 | 89 | 85 | 1.1 | 3 | 0.00477 |
| **GLYCEROPHOSPHOGLYCEROLS** | | | | | | | |
| | PG(39:4) | 13 | 113 | 22 | 5.2 | 198 | 1.90E-23 |
| | PG(ID_113) | 15 | 95 | 14 | 6.9 | 159 | 3.58E-22 |
| | PG(ID_349) | 7 | 30 | 27 | 1.1 | 39 | 3.14E-14 |
| **GLYCEROPHOSPHOINOSITOLS** | | | | | | | |
| | PI(ID_237) | 10 | 108 | 44 | 2.5 | 75 | 3.86E-18 |
| | PI(ID_62) | 13 | 121 | 23 | 5.2 | 87 | 5.08E-19 |
| **SPHINGOMYELINS** | | | | | | | |
| | SM(d34:1) | 8 | 156 | 23 | 6.7 | 124 | 4.80E-21 |
| | SM(d36:1) | 4 | 53 | 23 | 2.4 | 76 | 3.17E-18 |
